# Optogenetic control of kinesins -1, -2, -3 and dynein reveals their specific roles in vesicular transport

**DOI:** 10.1101/2023.04.18.537380

**Authors:** Sahil Nagpal, Samuel Wang, Karthikeyan Swaminathan, Florian Berger, Adam G. Hendricks

## Abstract

Each cargo in a cell employs a unique set of motor proteins for its transport. Often multiple types of kinesins are bound to the same cargo. It is puzzling why several types of motors are required for robust transport. To dissect the roles of each type of motor, we developed optogenetic inhibitors of kinesin-1, -2, -3 and dynein. This system allows us to control the activity of the endogenous set of motor proteins that are bound to intracellular cargoes. We examined the effect of optogenetic inhibition of kinesins-1, -2, and -3 and dynein on the transport of early endosomes, late endosomes, and lysosomes. While kinesin-1, kinesin-3, and dynein transport vesicles at all stages of endocytosis, kinesin-2 primarily drives late endosomes and lysosomes. In agreement with previous studies, sustained inhibition of either kinesins or dynein results in reduced motility in both directions. However, transient, optogenetic inhibition of kinesin-1 or dynein causes both early and late endosomes to move more processively by relieving competition with opposing motors. In contrast, optogenetic inhibition of kinesin-2 reduces the motility of late endosomes and lysosomes, and inhibition of kinesin-3 reduces the motility of endosomes and lysosomes. These results suggest that the directionality of transport is likely controlled through regulating kinesin-1 and dynein activity. On vesicles transported by several kinesin and dynein motors, motility can be directed by modulating the activity of a single type of motor on the cargo.

## Introduction

Kinesin and dynein drive the long-range transport of vesicular cargoes, mRNA, and proteins along microtubules, and their dysfunction often leads to neurodegenerative disease (1, 2). The activation and recruitment of motors are precisely coordinated to control the movement of specific cargoes and target them to different destinations in the cell (3). Multiple regulatory mechanisms direct cargoes to their destinations in the cell including scaffolding and adaptor proteins, microtubule-associated proteins, and mechanical interactions between different motors on the same cargo. In this study, we seek to mimic the endogenous mechanisms of regulation to better understand how perturbations in motor activity control the direction and processivity.

Conventional approaches to inhibit motor proteins, such as function-blocking antibodies (4), genetic inhibition (5), chemical-genetic inhibition (6), and dominant-negative expression (7) lack temporal control over regulation or are irreversible. Further, inhibition of either kinesins or dynein often stops motility in both directions (4, 5, 8). Optogenetics offers the potential to overcome these limitations and provide spatiotemporal control over motor activity (reviewed in (9)). Existing optogenetic applications to control motor activity have focused on recruiting exogenous motors to vesicular cargoes (10–12). While these systems provide insight into how the localization of organelles is affected by recruiting motors of different types to their surface, they are limited in their ability to probe the dynamics of the native sets of motors bound to intracellular cargoes.

Here, we developed optogenetic tools to reversibly inactivate endogenous kinesin-1, kinesin-2, kinesin-3, and cytoplasmic dynein. We used native autoinhibitory domains of kinesins and dynein to develop optogenetic inhibitors based on the LOVTRAP system (13). LOVTRAP employs a light-sensitive LOV2 protein, and a Zdark (or Zdk) protein that selectively binds to the dark-adapted conformational state of LOV2. We fused inhibitory domains for each motor to the Zdk protein. The LOV2 protein is anchored to the mitochondria membrane. In the absence of blue light (dark state), the inhibitory domain is sequestered on mitochondria. Upon blue light excitation, the Zdk protein is freed from the mitochondria due to conformational changes in LOV2, allowing the attached inhibitory domain to diffuse in the cytoplasm and interact with endogenous motor proteins (Figure 1 A, B, D and E). This LOV2 based trapping of inhibitory domains provides reversible control of motor activity with diffusion-limited activation kinetics. We used these optogenetic tools to dynamically control the localization of peptides that selectively inhibit kinesins 1-3 and dynein. We examined the effect of transiently inhibiting these motors on the motility of early endosomes, late endosomes and lysosomes to understand how teams of different motors work together to transport cargoes along microtubules. We find that optogenetic inhibition of kinesin-1 and dynein enhances transport by relieving competition between opposing motors, while inhibition of kinesin-2 and kinesin-3 reduces motility. Together, these results suggest that kinesin-1 and dynein control the direction of movement, while kinesin-2 and kinesin-3 aid in long-range transport.

**Fig. 1.**
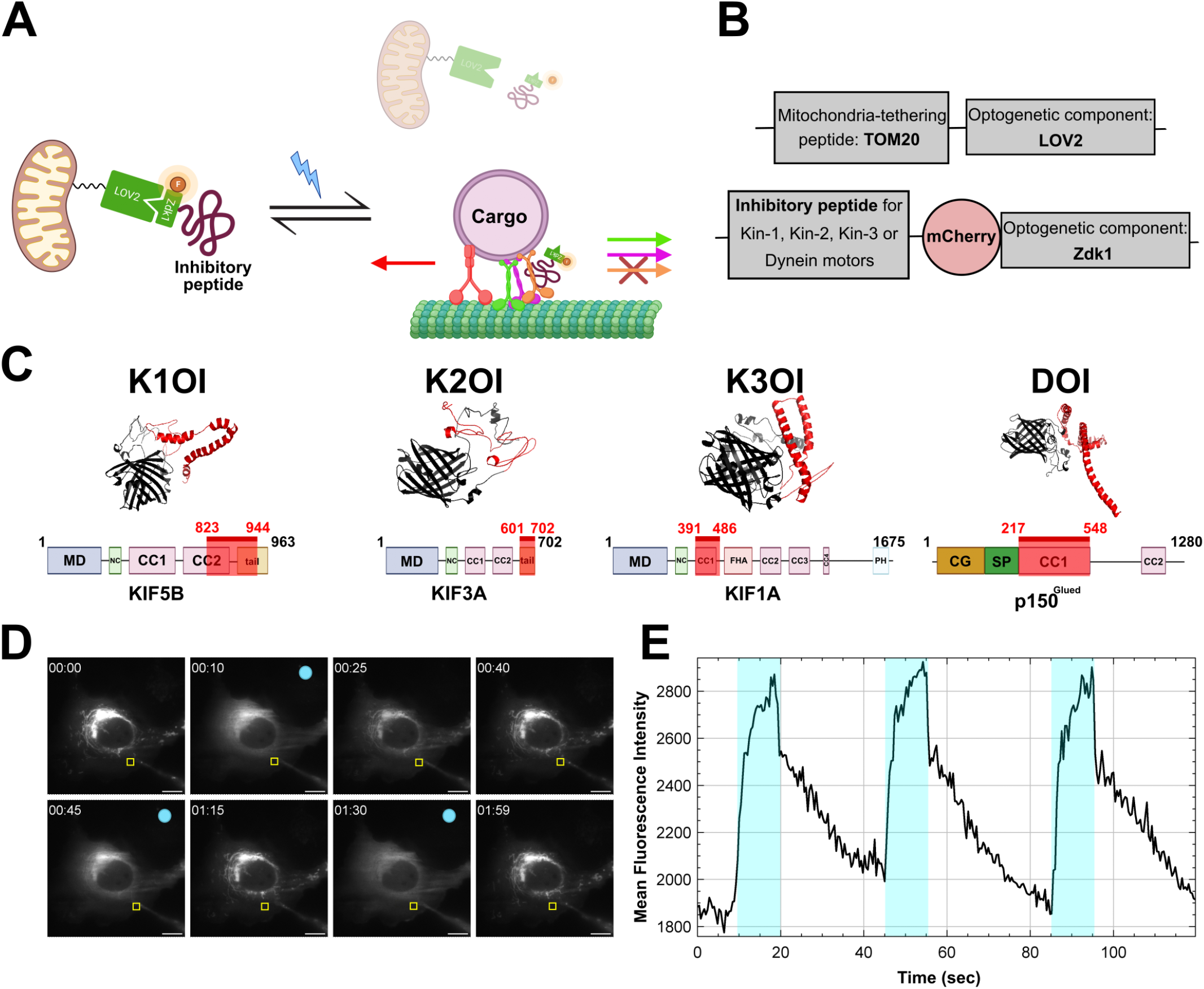
Using native autoinhibitory domains to build optogenetic inhibitors. (A) Schematic illustration of the mechanism of action for optogenetic inhibitors. In the dark-state, the inhibitory peptide is sequestered on mitochondria. Upon blue light illumination, it is released into the cytoplasm where it interacts with endogenous motor proteins. (B) Overview of plasmid constructs co-transfected for the optogenetic experiments. Plasmid construct on top encodes the protein that is tethered to the mitochondria and the bottom plasmid construct encodes the fluorophore containing inhibitory peptide. (C) Ab-initio based protein-prediction results from I-TASSER server (top) for different optogenetic inhibitors where the segment in red shows the cloned inhibitory peptide. Domain maps of different proteins used for creating optogenetic inhibitors (bottom) with the cloned segment highlighted in red. MD, motor domain; NC, neck coil; CC, coiled-coil; FHA, forkhead-associated; PH, pleckstrin homology; CG, CAP-Gly; SP, serine-proline rich region (D) Snapshots from a time-lapse imaging of a cell expressing the optogenetic constructs. Yellow box shows the cytoplasmic region used for generating the intensity trace shown in E. Scale bar 12µm. (E) Fluorescence intensity trace over time for the region of interest in cytoplasm for the cell shown in D, demonstrating the reversible aspect of optogenetic inhibitors.

## Results

### Optogenetic inhibitors of kinesin and dynein motors

Motor proteins adopt autoinhibitory states that allow them to remain inactive when not bound to a cargo, preventing unnecessary ATP hydrolysis and the congestion of microtubule tracks. Autoinhibition is often achieved by intra-molecular interactions between different domains of the motor protein ((14, 15)). We have exploited these interactions to design optogenetic tools that inhibit endogenous motor proteins in living cells upon illumination with blue light (Figure 1C).

When no cargo is bound to kinesin-1, it adopts a compact, autoinhibited conformation through a series of intramolecular interactions (16). In this autoinhibited state, the C-terminal globular tail domain is positioned near the enzymatically active heads (17). The highly-conserved QIAKPIRP amino acid sequence in the tail domain interacts with the Switch I region of the head domain and inhibits the initial microtubule-stimulated ADP release step of the motor (18). Peptides with a central QIAKPIRP sequence are capable of inhibiting kinesin-1 motility *in-vitro* (19). For the kinesin-1 optogenetic inhibitor (K1OI), we cloned the tail domain (amino acids 823-944) from the ubiquitous isoform of the kinesin heavy chain, KIF5B, onto the Zdk protein of the LOVTRAP system. However, the resulting protein enriched on microtubules due to the presence of basic residues N-terminus to the critical QIAKPIRP sequence (18), preventing its association with LOV2. Deletion of these basic residues (amino acids 904-918) reduced the affinity of K1OI to microtubules such that its localization could be controlled using LOVTRAP.

A similar autoinhibited state is observed in kinesin-2 (KIF3A/B) in the absence of cargo. However, in case of kinesin-2, the tail domain blocks the interaction of the motor with microtubules whereas the coiled-coil segment inhibits processive motility of the motor domain (14). We used the minimal inhibitory domain, amino acids 601 to 702 of KIF3A, to construct our kinesin-2 optogenetic inhibitor (K2OI).

In contrast to kinesin-1 and kinesin-2, autoinhibition in kinesin-3 motors is not dependent on folding of the protein, rather the coiled-coil 1 region is capable of directly interacting with the neck coil-coil, which prevents the dimer formation required for processive motility (20). To construct our kinesin-3 optogenetic inhibitor (K3OI), we used the coiled-coil 1 domain, along with the ß-finger domain of KIF1A (amino acids 391-486), both of which are essential for inhibition of motor activity (21). The amino acid sequences of these domains are homologous to KIF1B and KIF1C (Figure S1), suggesting it will interact with KIF1A, B, and C isoforms.

Cytoplasmic dynein 1 exists in an autoinhibited state in the absence of its binding partners. While recent studies have shed light on the mechanism of autoinhibition (15, 22), the picture remains incomplete. Thus, we adopted a different approach to optogenetically inhibit dynein motors. Instead of relying on a domain that is involved in an intramolecular interaction-based inhibition, we used used the residues of p150 that are required for the assembly of the the dynein-dynactin complex. Overexpression of the CC1 domain of p150 competes with endogenous p150 for binding with the intermediate chain, thereby disrupting functional dynein-dynactin complexes (23, 24). For the dynein optogenetic inhibitor (DOI), we cloned the CC1 domain of p150 onto the Zdk component of the optogenetic system, mimicking its overexpression during blue light illumination.

### Optogenetic inhibition of kinesin-1 and dynein enhances early endosome motility by relieving competition between opposing motors

Signalling molecules, macromolecules, and particles are transported in endosomes into and out of cells. Early endosomes are formed upon internalization of material from cell’s environment, and are characterised by the presence of specific membrane markers such as EEA1 and Rab5 proteins (25). Early endosomes move in short, fast runs interspersed with frequent pauses (26). Kinesin-1 (27), kinesin-2 (28), kinesin-3 (29, 30), kinesin-14 (27), and dynein (8) motors have all been reported to contribute to early endosome motility.

To understand the change in motion of cargoes upon inhibition, we characterised their motility over time using three parameters: the radius of gyration (Rg) is an indicator of the displacement, mean squared displacement (MSD) indicates processivity, and the running-mean velocity indicates the direction of movement (see methods). Positive velocities correspond to motility towards the cell periphery, while negative velocities correspond to inward movement.

We first examined cargo motility upon expressing just the inhibitory peptide without expressing the optogenetic component, which served as a non-optogenetic control to study the efficacy of the inhibitors. We observed a significant decrease in the Rg and processivity of early endosome in cells expressing the inhibitors in comparison to non-transfected cells, with the most profound drop in the case of dynein and kinesin-1 inhibition (Figure S2B,G). Concomitantly, there was an increase in stationary motility upon kinesin-1 inhibition, and more outward motility in the case of dynein inhibition (Figure S2H) with a shift from peripheral localization to juxtanuclear localization of early endosomes in all the inhibitory conditions (Figure S2F).

Optogenetic inhibition of different motors had varied effects on motility. Upon inhibition of kinesin-1 and dynein motors, we saw an overall increase in mean displacement (measured using the radius of gyration, Rg) and processivity (quantified using the mean-squared displacement, MSD) (Figure 2E,F). We grouped the endosomes based on their motility in the 20s period before the inhibition as outward (velocity*≥* 0.01 *µ*m/s), inward (velocity 0.01 *µ*m/s), or stationary (*≤*0.01 *µ*m/s < velocity < -0.01 *µ*m/s) and analyzed cargoes that we could track over the entire time-lapse movie. We observe that upon kinesin-1 inhibition, the velocity of cargoes moving towards the cell periphery becomes more variable, suggesting that kinesin-1 inhibition increases the competition between opposing motors leading to an enhanced bidirectional motility. Stationary cargoes move inward upon kinesin-1 inhibition. Dynein inhibition leads to a directional switch where minus-end directed cargoes reverse and move to the cell periphery (Figure 2D). This data suggests an indirect effect of motor inhibition on net motility, whereby inhibiting kinesin-1 and dynein motors relieves the competition between opposing motors. Kinesin-2 is not a primary motor on early endosomes ((30, 31)). Accordingly, early endosomes were not strongly affected by kinesin-2 optogenetic inhibition (Figure 2E), consistent with long-term overexpression of the inhibitory peptide (Figure S2). While inhibiting kinesin-1 and dynein enhanced early endosome motility, inhibiting kinesin-3 motors resulted in decreased displacement and processivity (Figure 2E,F) and a shift towards inward movement (Figure 2D), suggesting kinesin-1 and kinesin-3 have different roles in driving the motility of early endosomes.

**Fig. 2.**
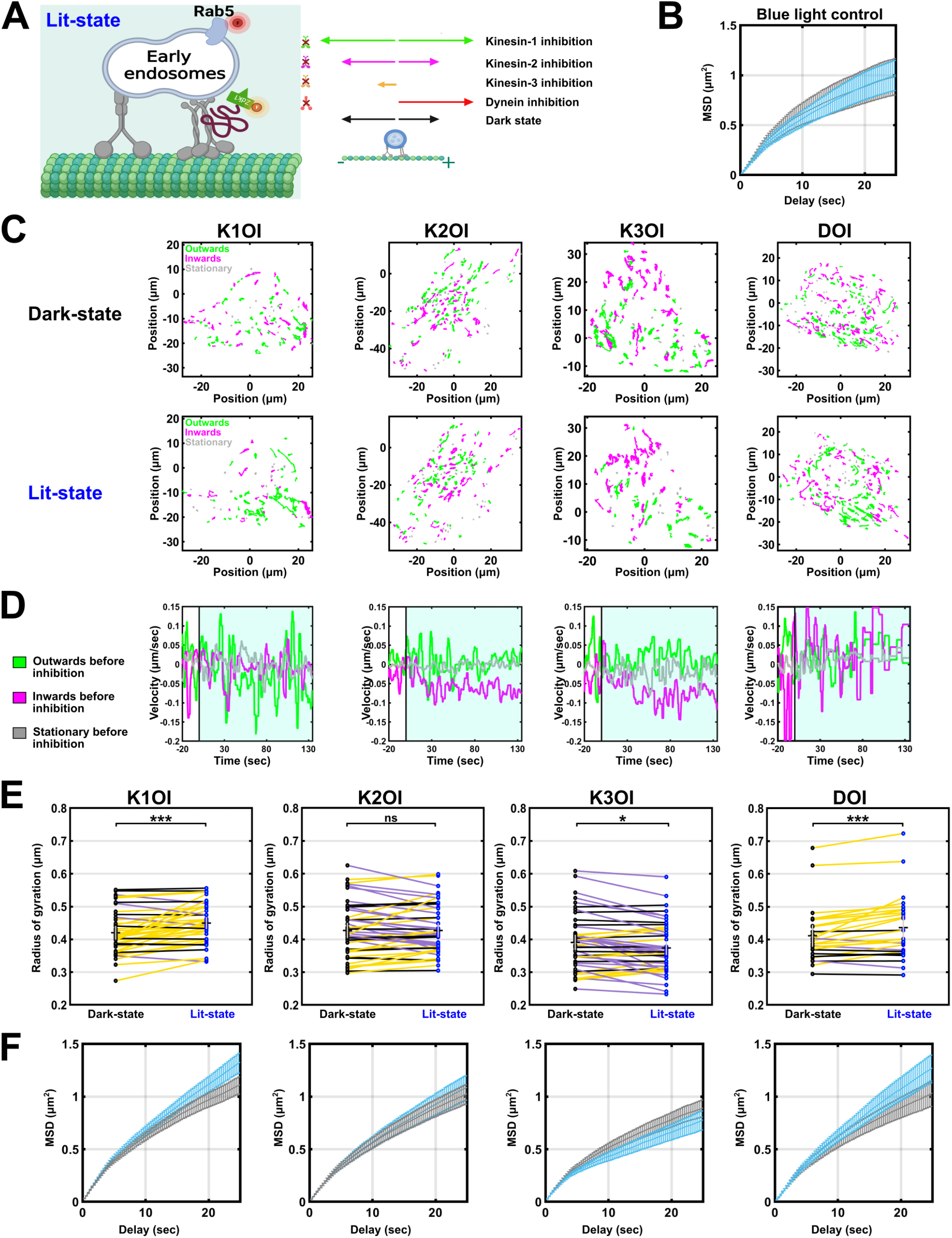
Early endosome motility is modulated by optogenetic inactivation of kinesin-1 & 3 and dynein motors. (A) Scheme showing the inhibition of motors that are driving early endosomes in the lit-state (left), with the inhibitory peptide labelled with an orange fluorophore while the early endosome marker, Rab5, is labelled with a far-red fluorophore. On the right is the summary of change in **Fig. 2**. motility upon inhibition of different transport motors, shown by differently colored arrows where the length of the arrow indicates the run length of the cargo. (B) The mean-squared displacement (MSD) plot of early endosomes in untransfected U2OS cells, that do not express optogenetic inhibitors, without and with blue light illumination, shown in black and blue respectively (mean ± SEM). Each cell was first imaged without shining any blue light, and then with blue light illumination. This blue light control shows that blue light itself does not affect the motility of early endosomes. (C) Polar plot projections of early endosomes trajectories from time-lapse images, centered around the cell nucleus, showing the directionality of Rab-5 enriched endosomes in a U2OS cell under dark-state (upper panel) and lit-state (bottom panel) conditions. The four panels correspond to cells that were transiently transfected with different optogenetic inhibitors. The net directionality was categorized as inwards (magenta), outwards (green) or stationary (gray) based on Rg values, and rho values in the first and the last points of the trajectories. (D) Plot shows the changes in average velocity for all the trajectories in a cell (corresponding to the cell shown in C) upon blue light illumination. For velocity analysis, average velocity was first categorised into three types, namely, positive velocity, negative velocity, and neutral velocity. It was then normalized to the average velocity in the time window just before inhibition, allowing us to compare changes at the time of inhibition. The color scheme is also based on the average velocity of the trajectories right before the inhibition, where green represents positive average velocity before inhibition, magenta represents negative average velocity before inhibition, and gray represents stationary vesicles that were not moving before inhibition. (E and F) Radius of gyration (Rg) and MSD plots for motility of early endosomes upon optogenetic inhibition of different motors. Each dot in the Rg plot indicates a cell, with a line connecting the same cell under the two conditions. A yellow line indicates an increase in Rg, whereas a purple line indicates a decrease, and a black line indicates no change. Black horizontal line shows mean while vertical gray line indicates SEM. For the MSD plot, dark-state and lit-state are shown in gray and blue respectively (mean ± SEM). The number of cells, trajectories and experiments used for the plots are as follows:-K1OI: 41 cells, 7713 trajectories over 5 experiments; K2OI: 48 cells, 8560 trajectories over 3 experiments; K3OI: 47 cells, 8976 trajectories over 5 experiments; DOI: 26 cells, 4859 trajectories over 3 experiments. Statistical significance for Rg analysis was done using Wilcoxon signed-rank test and asterisks indicate significance as follows: *∗∗∗* for p *≤* 0.001, *∗∗* for p *≤* 0.01, *∗* for p *≤* 0.05.

### Late endosome motility is enhanced with optogenetic inhibition of kinesin-1 and dynein, and disrupted by inhibition of kinesin-2 & -3

As endosomes progress in the endocytic pathway, they undergo morphological and biochemical changes as they mature into late endosomes (the process is reviewed in detail in (32)). Maturation also involves luminal acidification (with pH dropping from around 6.5 to 5.5 (33)) and movement towards the perinuclear region with more frequent directional switches (34). This transition is accompanied by a switch from Rab5 to Rab7, a dynamic process initiated by recruitment of Rab7-GEF by Rab5-GTP and subsequent release of Rab5-GEF, preventing further Rab5 activation (35). Rab7 recruits other adaptor proteins such as RILP and FYCO, that enables interaction with dynein and kinesin motors respectively, thus allowing late endosomes to move bidirectionally (36, 37).

Upon overexpression of different motor inhibitors, we again observed the most significant change in mean Rg and MSD in the case of kinesin-1 and dynein inhibition. However, unlike early endosomes, inhibition of kinesin-2 showed a more pronounced effect on the motility of late endosomes (Figure S3B,G). Apart from dampened motility, we also observed clustering of vesicles, which is in concert with a previous study that showed clustering of late endosomes upon kinesin-2 inhibition (31). For all inhibitory conditions, we observed an increase in stationary runs and a slight shift from perinuclear enrichment of vesicles to increased juxtanuclear and peripheral enrichment (Figure S3F,H). Therefore, sustained inhibition of motors impairs the transport kinetics of these vesicles, steering them to improper locations within the cell.

Rab7-positive vesicles responded similarly to Rab-5-positive vesicles to optogenetic motor inhibition. The radius of gyration of late endosomes increased upon transient inhibition of kinesin-1 and dynein motors, indicative of relieving competition between opposing motors (Figure 3E). The effect on processivity is less apparent (Figure 3F), because longer trajectories are weighed more heavily in the MSD analysis while each trajectory is weighed equally in the Rg calculation, suggesting that kinesin-1 and dynein inhibition has a stronger effect on short trajectories. Upon kinesin-1 inhibition, we observe that the vesicles that were moving towards the plus end switch to dynein-driven minus ended motility upon blue light illumination. When dynein is inhibited, inward trajectories switch to outward movement while outward trajectories continue to move outward (Figure 3D). Kinesin-2 and kinesin-3 inhibition results in a clear decrease in displacement and processivity, indicating that these motors drive long-range transport of late endosomes (30, 31).

**Fig. 3.**
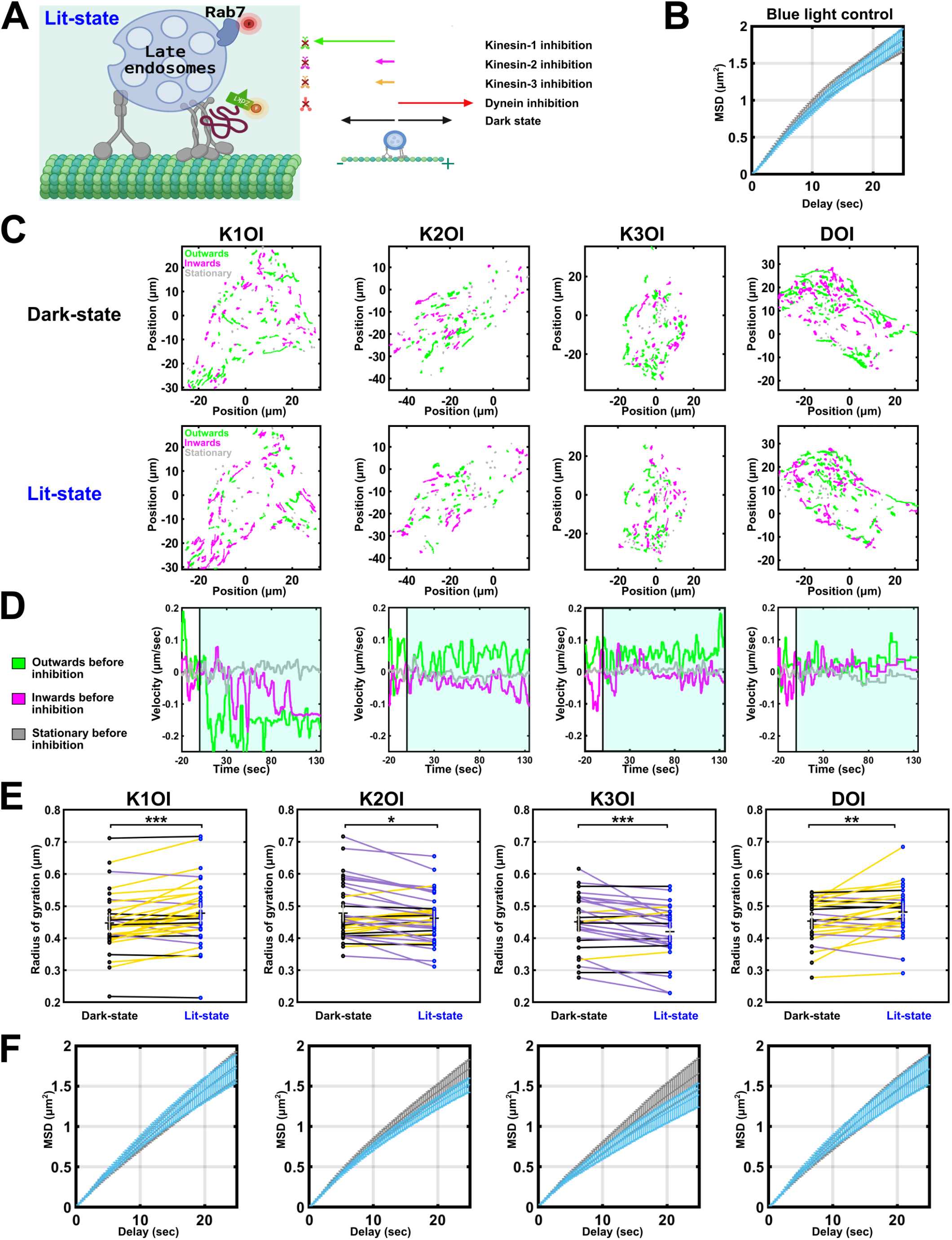
Late endosome motility is enhanced by optogenetic inhibition of kinesin-1 and dynein, and reduced by optogenetic inhibition of kinesin -2 & -3. (A) Scheme showing the inhibition of motors that are driving late endosomes in the lit-state (left), with the inhibitory peptide labelled with an orange fluorophore while late endosome marker, Rab7, is labelled with a far-red fluorophore. **Fig. 3**. On the right is the summary of change in motility upon inhibition of different transport motors, shown by differently colored arrows where the length of the arrow indicates the run length of the cargo. (B) The mean-squared displacement (MSD) plot of late endosomes in untransfected U2OS cells, that do not express optogenetic inhibitors, without and with blue light illumination, shown in black and blue respectively (mean ± SEM). Each cell was first imaged without shining any blue light, and then with blue light illumination. This blue light control shows that blue light itself does not affect the motility of late endosomes. (C) Polar plot projections of late endosomes trajectories from time-lapse images, centered around the cell nucleus, showing the directionality of Rab-7 enriched endosomes in a U2OS cell under dark-state (upper panel) and lit-state (bottom panel) conditions. The four panels correspond to cells that were transiently transfected with different optogenetic inhibitors. The net directionality was categorized as inwards (magenta), outwards (green) or stationary (gray) based on Rg values, and rho values in the first and the last points of the trajectories. (D) Plot shows the changes in average velocity for all the trajectories in a cell (corresponding to the cell shown in C) upon blue light illumination. For velocity analysis, average velocity was first categorised into three types, namely, positive velocity, negative velocity, and neutral velocity. It was then normalized to the average velocity in the time window just before inhibition, allowing us to compare changes at the time of inhibition. The color scheme is also based on the average velocity of the trajectories right before the inhibition, where green represents positive average velocity before inhibition, magenta represents negative average velocity before inhibition, and gray represents stationary vesicles that were not moving before inhibition. (E and F) Radius of gyration (Rg) and MSD plots for motility of late endosomes upon optogenetic inhibition of different motors. Each dot in the Rg plot indicates a cell, with a line connecting the same cell under the two conditions. A yellow line indicates an increase in Rg, whereas a purple line indicates a decrease, and a black line indicates no change. Black horizontal line shows mean while vertical gray line indicates SEM. For the MSD plot, dark-state and lit-state are shown in gray and blue respectively (mean ± SEM). The number of cells, trajectories and experiments used for the plots are as follows:-K1OI: 34 cells, 5638 trajectories over 4 experiments; K2OI: 37 cells, 6189 trajectories over 3 experiments; K3OI: 28 cells, 4473 trajectories over 3 experiments; DOI: 27 cells, 4512 trajectories over 3 experiments. Statistical significance for Rg analysis was done using Wilcoxon signed-rank test and asterisks indicate significance as follows: *∗∗∗* for p *≤* 0.001, *∗∗* for p *≤* 0.01, *∗* for p *≤* 0.05.

### Lysosome motility is enhanced upon optogenetic inhibition of dynein and supressed by inhibition of kinesins -1, -2, & -3

The majority of late endosomes fuse with lysosomes, generating a hybrid organelle called an endolysosome that provides a controlled acidic environment for the degradation of the endocytosed macromolecules. These endolysosomes further mature to form dense lysosomes, which is the final compartment of the endocytic pathway (38). Lysosomes have a characteristically low pH of around 4.5, owing to the presence of degradative acid hydrolases (33, 38). Lysosome motility involves kinesin-1, -2 and -3 for anterograde transport and dynein for retrograde transport (reviewed in (39)).

Overexpression of different motor inhibitors showed markedly different effects on lysosome motility compared to Rab5- and Rab7-enriched vesicles. Kinesin-3 and dynein inhibition reduced the mean Rg and MSD with an almost two and three times drop in fraction of high Rg lysosomes respectively (Figure S4B,G). Furthermore, there was an increase in the fraction of stationary lysosomes upon kinesin-3 and dynein inhibition. Kinesin-1 and kinesin-2 inhibition did not significantly affect the mean Rg or directionality of the cargoes, however, there was a decrease in processivity, as shown by the MSD. Similar to late endosomes, we observe a shift from perinuclear localization of lysosomes to a more peripheral localization for all inhibitors, with the most significant change in case of kinesin-1 inhibition (Figure S4F).

Lysosomes also responded differently to optogenetic inhibition of motors compared to early and late endosomes (Figure 4 and S5). For lysosomes, kinesin-1 inhibition resulted in reduced motility (Figure 4E,F), in contrast to early and late endosomes (Figures 2E,F, 3E,F). The cargo velocities provide insight into the effect of the inhibitor, where both inward and outward motility becomes stationary upon inhibition of kinesin-1 with no strong directional switches. These results suggest kinesin-1 motors play a different role in lysosome motility compared to endosomes (Figure 4D). Kinesin-2 and kinesin-3 inhibition led to a decrease in mean Rg and MSD with a clear shift towards minus-ended motility in the lit-state (Figure 4 D, E and F). For dynein inhibition, we see enhanced motility with a directional switch towards the plus end of the microtubules, although the effect is weaker than observed for early and late endosomes. Interestingly, lysosomes that were not motile in the dark-state started moving towards the cell periphery upon dynein inhibition (Figure 4D).

**Fig. 4.**
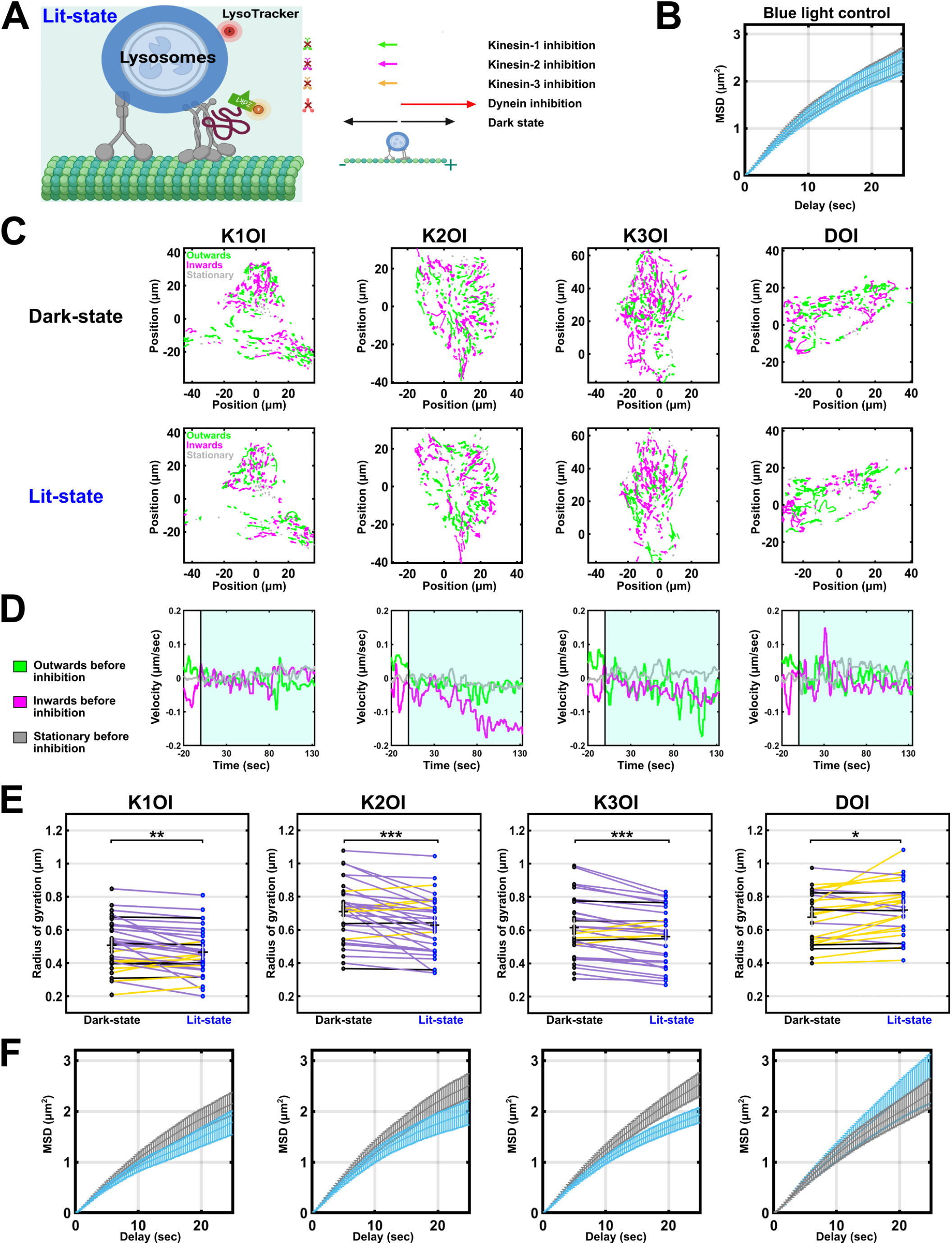
Lysosome motility is reduced by acute inhibition of kinesin motors, and enhanced upon acute inhibition of dynein. (A) Scheme showing the inhibition of motors that are driving lysosomes in the lit-state (left), with the inhibitory peptide labelled with an orange fluorophore while the lysosome is labelled with far-red LysoTracker. On the right is the summary of change in motility upon **Fig. 4**. inhibition of different transport motors, shown by differently colored arrows where the length of the arrow indicates the run length of the cargo. (B) The mean-squared displacement (MSD) plot of lysosomes in untransfected U2OS cells, that do not express optogenetic inhibitors, without and with blue light illumination, shown in black and blue respectively (mean ± SEM). Each cell was first imaged without shining any blue light, and then with blue light illumination. This blue light control shows that blue light itself does not affect the motility of lysosomes. (C) Polar plot projections of lysosomes trajectories from time-lapse images, centered around the cell nucleus, showing their directionality in a U2OS cell under dark-state (upper panel) and lit-state (bottom panel) conditions. The four panels correspond to cells that were transiently transfected with different optogenetic inhibitors. The net directionality was categorized as inwards (magenta), outwards (green) or stationary (gray) based on Rg values, and rho values in the first and the last points of the trajectories. (D) Plot shows the changes in average velocity for all the trajectories in a cell (corresponding to the cell shown in C) upon blue light illumination. For velocity analysis, average velocity was first categorised into three types, namely, positive velocity, negative velocity, and neutral velocity. It was then normalized to the average velocity in the time window just before inhibition, allowing us to compare changes at the time of inhibition. The color scheme is also based on the average velocity of the trajectories right before the inhibition, where green represents positive average velocity before inhibition, magenta represents negative average velocity before inhibition, and gray represents stationary vesicles that were not moving before inhibition. (E and F) Radius of gyration (Rg) and MSD plots for motility of lysosomes upon optogenetic inhibition of different motors. Each dot in the Rg plot indicates a cell, with a line connecting the same cell under the two conditions. A yellow line indicates an increase in Rg, whereas a purple line indicates a decrease, and a black line indicates no change. Black horizontal line shows mean while vertical gray line indicates SEM. For the MSD plot, dark-state and lit-state are shown in gray and blue respectively (mean ± SEM). The number of cells, trajectories and experiments used for the plots are as follows:-K1OI: 34 cells, 3791 trajectories over 4 experiments; K2OI: 34 cells, 3843 trajectories over 4 experiments; K3OI: 34 cells, 3798 trajectories over 5 experiments; DOI: 27 cells, 3114 trajectories over 3 experiments. Statistical significance for Rg analysis was done using Wilcoxon signed-rank test and asterisks indicate significance as follows: *∗ ∗ ∗* for p *≤* 0.001, *∗∗* for p *≤* 0.01, *∗* for p *≤* 0.05.

### Mathematical modeling reveals the estimated fraction of inhibited motors

To gain more insight into how the mechanochemical properties of different motors determines their roles in vesicle transport, we tested the effect of inhibiting specific motors in a mathematical model of transport by kinesins and dynein. We extended the mathematical model developed by Müller et al. (40) to simulate the motility of a cargo driven by any number of different types of motors (see Methods). We estimated motor parameters such as binding rate, detachment rate, detachment force, stall force, forward and backward velocities based on previous single-molecule experiments and published models, following closely the analysis in (41) (see methods). To simulate different vesicle populations, we used estimates of the number of motors on each of these cargoes from previous single-molecule fluorescence and quantitative photobleaching experiments (42). Using these parameters, the model captured the motility characteristics early endosomes, late endosomes, and lysosomes. We then mimicked the effect of optogenetic inhibition of the motors in the model by decreasing the binding rates of motors.

Simulated trajectories show similar changes in displacement and directionality in response to motor inhibitions as observed experimentally (Figure 5A). We used the model to estimate the degree to which optogenetic inhibitors decreased the activity of different motors in our experiments. We estimate the optogenetic inhibitors developed here reduced the binding rate by *∼*40-60% for kinesin-1, 30-40% for kinesin-2, 30-50% for kinesin-3, and 25-50% for dynein depending on the cargo (Fig. 5A). We also examined the change in directionality of simulated trajectories upon inhibition of different motors (Figure 5B). We observed a similar trend in change of directionality as our experimental data. For lysosomes, the change in directionality from optogenetic inhibition in the experiment was greater than the estimated change from the model, indicating that there could be cross-talk between the motors when several motor types are active at the same time, where inhibiting one type of kinesin also inhibits other types of kinesins bound to the same cargo, therefore creating a stronger response.

**Fig. 5.**
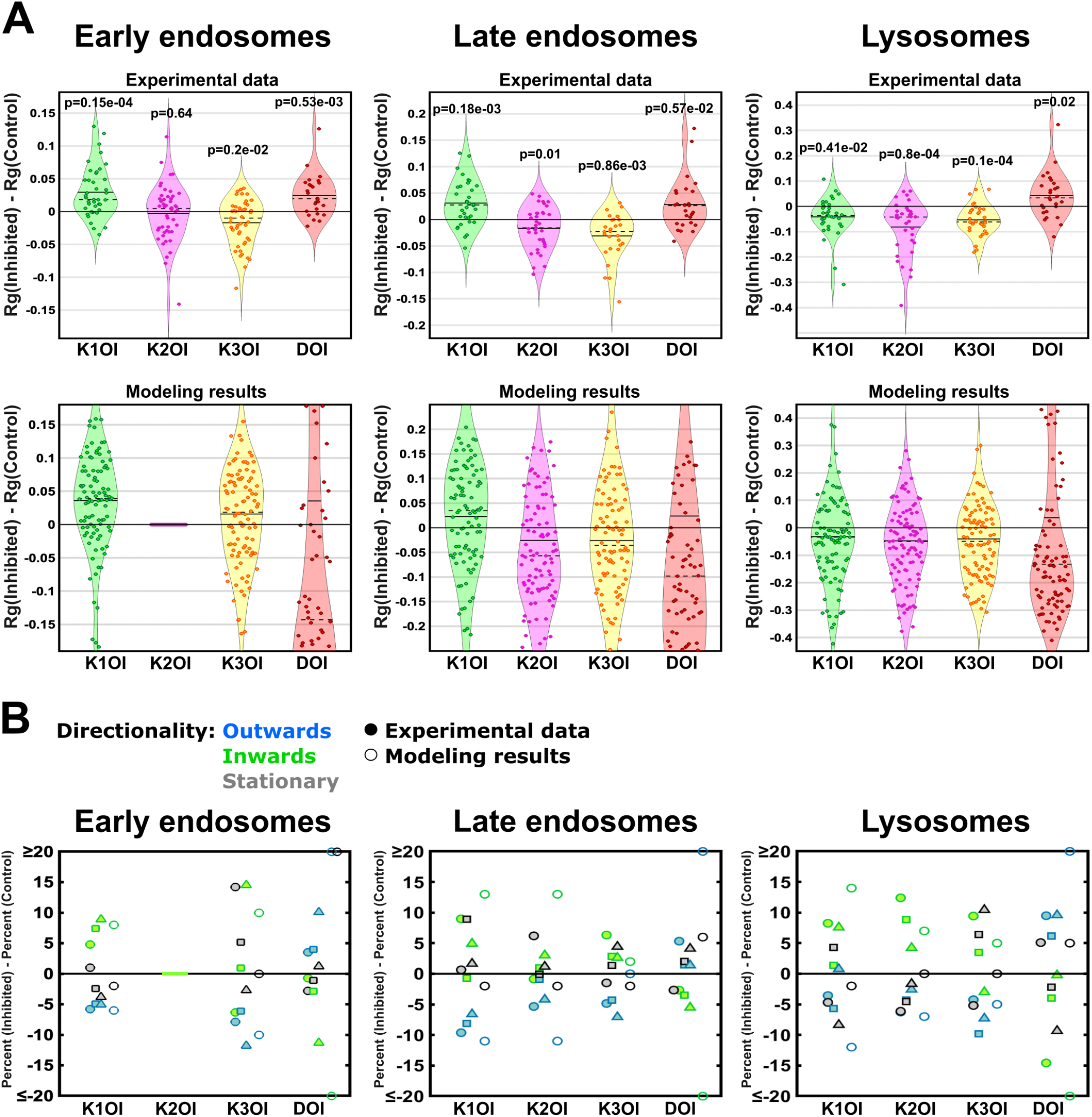
Mathematical modeling indicates the unique mechanochemical properties of each motor determines its role in transport. (A) Plots for difference in Rg upon optogenetic inhibition of different motors for early endosomes, late endosomes and lysosomes. Solid horizontal line indicates mean, and dashed horizontal line indicates median. Upper panel summarizes the Rg results reported in figures above and the lower panel indicates the normalized Rg values obtained from the modeled trajectories under different inhibitory conditions. The simulated trajectories exhibit a broader distribution as results for each simulated trajectory are plotted, compared to the mean of the trajectories for each cell in the experimental data. Binding rates are estimated to be reduced by *∼*40-60% for K1OI, 30-40% for K2OI, 30-50% for K3OI, and 25-50% for DOI depending on the cargo. P-values were calculated from student’s t-test. (B) Plots indicating changes in directionality upon inhibition of different motors for early endosomes, late endosomes and lysosomes. Trajectories were categorized into outwards, inwards or stationary based on the difference in positions for end and start points of the trajectory. Filled circles, triangles, and squares indicate experimental data for three different cells, and empty circles indicate the modeling results.

## Discussion

Cargo transport is regulated through multiple mechanisms including through the cytoskeletal tracks, motor adaptor and scaffolding proteins, and mechanical interactions between motors on the same cargo. To examine how modulating the activity of cargo-bound motors governs their motility, we developed optogenetic inhibitors that target the endogenous kinesins and dynein carrying different cargoes.

Through using optogenetics to control kinesin and dynein activity, we addressed the following questions:

### What motors contribute to the motility of different endocytic cargoes?

While the types of motors associated with different vesicle populations have been characterized using biochemical approaches and live cell imaging (30, 42), determining which motors are active has been challenging as long-term inhibition via dominant negative expression or genetic manipulation often results in reduced transport in both directions (4, 5, 8). Through comparing transport in the same cell in control and under acute inhibition of specific motors, we could map which motors contribute to the motility of early endosomes, late endosomes, and lysosomes (Figure 6A). We find that early endosomes are transported by kinesin-1, and -3, and dynein. Kinesin-1, -2, and -3, and dynein all contribute to late endosome and lysosome motility (Fig. 6A).

**Fig. 6.**
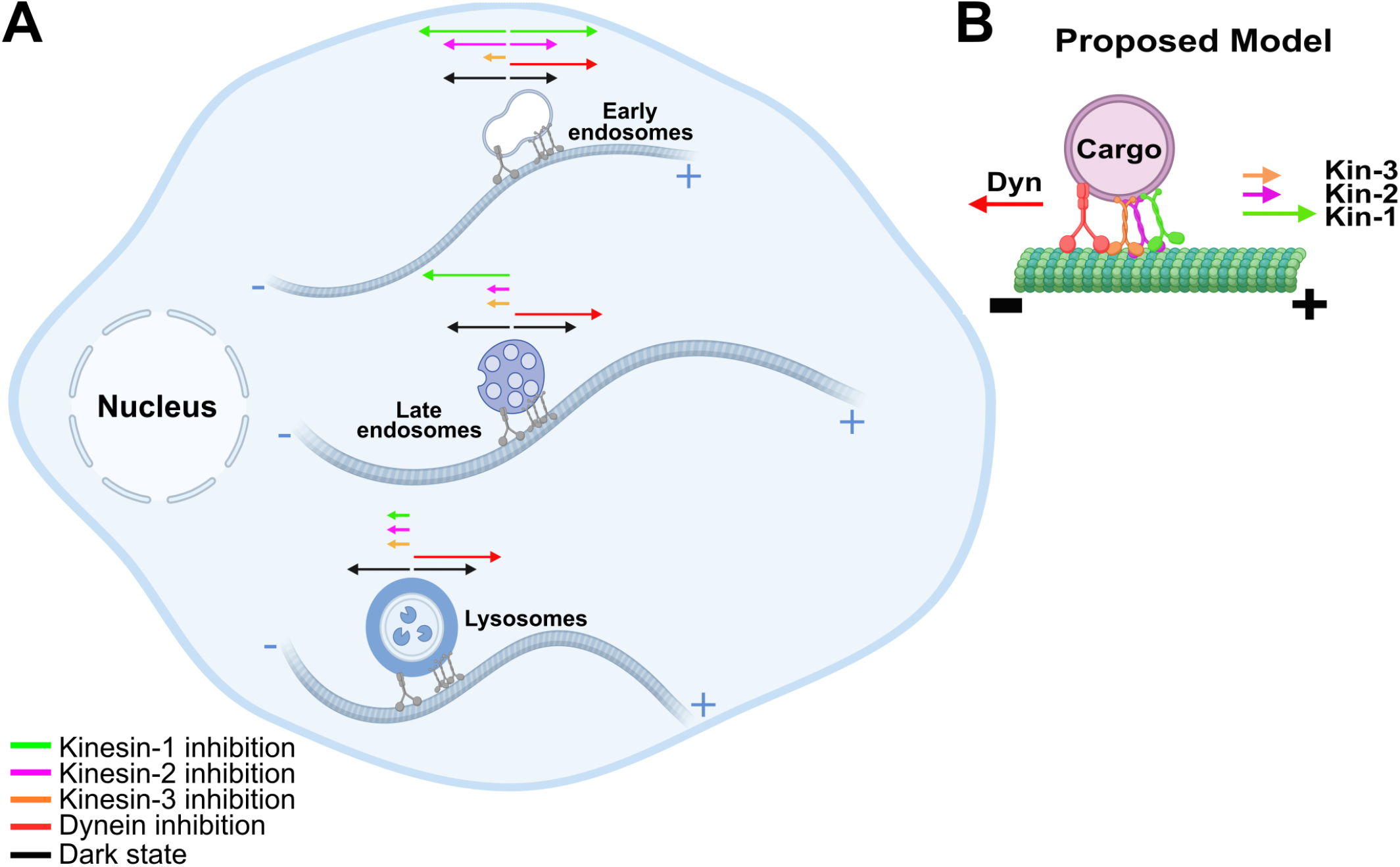
Different endocytic vesicles have varying responses to optogenetic inactivation of kinesins and dynein. (A) Schematic for transport of different endocytic vesicles based on optogenetic inhibition of different motor proteins. The length of the arrow indicates the change in motility of the cargo upon optogenetic inhibition. (B) Proposed model where cargo motility is more sensitive to the activity of kinesin-1 and dynein, while kinesin-2 and -3 may play a more supportive role to direct the cargo to its correct destination. The length of the arrow indicates the sensitivity of the respective motor in determining the net motility of the cargo.

### Why are multiple types of motors bound to the same cargo?

Kinesins -1, -2, and -3 are all processive and drive transport towards microtubule plus ends. Yet, many intracellular cargoes are bound by multiple types of kinesins and dynein (30, 42, 43). What advantage is provided by multiple kinesin types acting on the same cargo? Our results suggest each motor type has distinct roles in cargo transport. Optogenetic inhibition of kinesin-1 and dynein results in longer trajectories and a switch in the direction of movement (Fig. 5,6). Thus, kinesin-1 and dynein likely play regulatory roles to determine the direction of motility. In contrast, optogenetic inhibition of kinesin-2 and -3 results in reduced displacement (Fig. 5A) and more stationary trajectories (Fig. 5B), suggesting that kinesin-2 and -3 act as long-range haulers. Optogenetically inhibiting multiple kinesins at the same time showed a more prominent directional switch with the majority of cargoes shifting to dynein-driven motility (Figure S6), indicating that multiple types of kinesins work together to transport cargoes (44). Interestingly, microtubule-associated proteins (MAPs) like tau and MAP7 preferentially target kinesin-1 (43, 45), which is sensitive to tubulin lattice spacing (46). Kinesin-1 and -3 prefer microtubules marked with different set of post-translational modifications (47, 48). Further, multiple mechanisms are required to fully activate kinesin-1 (16, 49). Together, these results suggest that many mechanisms that regulate intracellular transport might specifically target kinesin-1, where kinesin-2 and -3 could aid in long-range transport (Figure 6B).

We found that sustained and acute inhibitions had differential effects on cargo motility. While overexpression of inhibitors often led to a halt in motility with more stationary runs, optogenetic inhibition upregulated cargo transport in some cases of motor inhibition. This indirect effect of acute inhibition suggests close coupling between different motors that are on a cargo, whereby motors can directly activate and influence each other. This cross-talk between opposite polarity motors has been previously established in other studies where it was shown that the slowest teams of motors mechanically communicate with other motors through the membrane of the cargo to drive transport (50, 51). Based on our results, we hypothesize that selectively inhibiting kinesin-1 and dynein motors in the case of early and late endosomes relieves competition with opposing motors. Turning off kinesin-1 or dynein might enable more processive motors, such as kinesin-3, to take over transport. This hypothesis is in line with computational and *in-vitro* studies that suggest that the number of engaged motors governs motility (52–54).

To summarize, we developed optogenetic inhibitors of kinesin-1, -2, -3, and dynein motors to mimic regulation by motor adaptors and effectors in the cell. We examined the change in motility of different endocytic vesicle populations upon acute, optogenetic inhibition. Acute inhibition has different effects on motility compared to long-term inhibition using dominant negatives. Short-term inhibition with optogenetics gave a glimpse of complex motor dynamics on these cargoes, where kinesin-1 and dynein inhibition leads to enhanced motility, owing to strong directional switches to the opposite direction, whereas kinesin-2 and -3 inhibition results in reduced motility. We propose that the activity of kinesin-1 and dynein motors determines the net direction of movement while kinesin-2 and -3 aid in long-range transport. In conclusion, different vesicles of the endocytic pathway rely on unique sets of transport motors, where modulating the activity of a single type of motor differentially affects the overall motility of the cargo, indicating an intricate underlying interplay of motors to drive the vesicle to its right destination.

## Materials and Methods

### DNA Constructs and cloning

The optogenetic module was derived from plasmids pTriEx-NTOM20-LOV2 and pTriEx-mCherry-Zdk1 (Addgene plasmid #81009 and #81057 respectively), gifts from Klaus Hahn. Markers for endosomes were derived from plasmids iRFP-FRB-Rab5 and iRFP-FRB-Rab7 (Addgene plasmid #51612 and #51613 respectively), gifts from Tamas Balla. Dominant negative constructs for KIF5B, KIF3A and dynein were kindly gifted by Erika Holzbaur. Dominant negative construct for kinesin-3 was constructed from plasmid pBa-FRB-3myc-KIF1A tail 391-1698, a gift from Gary Banker and Marvin Bentley (Addgene plasmid # 64286).

A peptide linker - GSGGSGSGGT - was added to the inhibitory peptide constructs before fusing them with mCherry-Zdk1. Optogenetics constructs were created with the overlap extension PCR cloning method (55) and the primers were designed on an online platform for restriction-free cloning (56). Briefly, an amplicon containing the fragment of interest from the donor plasmid and flanked by a nucleotide sequence that overlaps with the insertion site of the recipient plasmid was generated from the primary PCR step. This amplicon was gel extracted and was used as an oversized primer for the secondary PCR reaction. The recombinant plasmid obtained at the end of secondary PCR reaction was incubated with Dpn1 enzyme, followed by its transformation into bacteria. Finally, colony PCR was done to screen the colonies and positive clones were confirmed by Sanger sequencing.

### Cell culture and transfection

U-2 OS and COS-7 cells were obtained from American Type Culture Collection, Manassas, VA. All cell lines were routinely screened for mycoplasma contamination. Cells were grown on 100mm polystyrene cell culture dishes (Gibco, Thermo Fisher Scientific, Waltham, MA) for maintenance and on 35mm glass-bottom dishes - No. 1.0 coverslip (MatTek Corporation, Ashland, MA) for imaging, with DMEM basal media (Gibco), supplemented with 10% (v/v) FBS (Gibco), 1% (v/v) Glutamax (Gibco) and Penstrep(Gibco). Cells were passaged using 0.25% trypsin-EDTA (Gibco) and PBS (Wisent, St-Jean-Baptiste, QC, Canada) and incubated at 37°C with 5% CO2 for 24h prior to transient transfection. Cells were transiently transfected with ~400ng plasmid DNA prepared in OPTI-MEM Reduced Serum (Gibco), using Lipofectamine LTX with Plus-Reagent (Invitrogen, Thermo Fisher Scientific), according to the manufacturer’s instructions. Cells were incubated with transfection solution for four hours before replacement with fresh DMEM media. For staining lysosomes, cells were treated with 50nM Lysotracker Deep Red (Invitrogen) for 10 minutes in complete DMEM media.

### Microscopy and imaging

Live-cell imaging was performed on a TIRF set-up built on an Eclipse Ti-E inverted microscope (Nikon, Melville, NY) with an attached EMCCD camera (iXon U897, Andor Technology, South Windsor, CT), 1.49 numerical aperture oil-immersion 100x objective and 100 mW diode lasers (Coherent). Time-lapse movies were taken using 561nm laser (1 mW) to excite the mCherry fluorophore, 647nm laser (1 mW) to excite the endosome marker, and 488nm laser (1 mW) for blue-light illumination, along with appropriate emission filter sets.

Cells were imaged at 37°C in Leibovitz’s L-15 media (Gibco), supplemented with 10% (v/v) FBS. For overexpression studies, cells were co-transfected overnight with dominant-negative constructs at a concentration of 200ng of DNA per dish and endosome marker constructs at 50ng of DNA per dish. Endosomes were imaged for 120s, with exposure set to 300ms per frame. For optogenetic studies, cells were co-transfected overnight with pTriEx-NTOM20-LOV2 plasmid at a concentration of 300ng of DNA per dish, inhibitor construct at 50ng of DNA per dish (with the exception of K1OI, which was at 15ng of DNA per dish) and endosome marker constructs at 50ng of DNA per dish. mCherry imaging was done at an exposure of 800ms per frame for 45s, with a short pulse of blue-light illumination for 5s. Endosomes were imaged for 120s in the dark-state followed by 135s period in the lit-state (out of which, initial 15s period was assumed to be the inactivation period, where the inhibitors diffuse around the cytoplasm and interact with motors to deactivate them, and was not considered for MSD & Rg analysis), with exposure set to 300ms per frame.

### Tracking of cargoes

Endosomes were tracked with the ImajeJ plugin TrackMate (57), with the Laplacian of Gaussian filter, and sub-pixel localization for cargo detection, and the simple Linear Assignment Problem tracker that generates the trajectories. Based on the known differences in maximal velocity for different endosomes, tracking parameters were set to linking distance (*µ*m):gap-closing maximum distance (*µ*m) :gap-closing maximum frame gap of 1:1:1 for Rab5 and Rab7 vesicles and 1.5:1.5:1 for lysosomes. However, changing these parameters did not significantly affect the analysis (Fig. S7).

### Analysis of motion

Using Trackmate output, X and Y coordinates, and time duration of each cargo trajectory in a cell was obtained. 2D position of individual trajectories was calculated by taking the square root of the sum of the squares of the X and Y coordinates of the trajectory with respect to the origin of the trajectory. This position data of trajectories was then used for further analysis.

Radius of gyration (Rg) is the radius of a circle that can be drawn around the trajectory, in such a way that it encompasses half of the points in the trajectory, thus parameterizing the size of the trajectory. It is indicative of the distance covered by the motor-driven cargo, indirectly estimating the run length of the motor. Radius of gyration is a scalar quantity (thus directionless), calculated using the following formula:

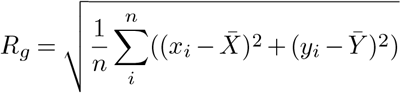

where n is the total number of detected points in the trajectory at consecutive time frames, x_i_ and y_i_ are the X and Y coordinates of the trajectory point at i time point, and 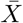 and 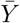 are the mean position of all the points in the trajectory. In our study, we took the mean of calculated Rg values for all the trajectories in a cell.

MSD is the average of the squared (therefore directionless) displacement of the cargo from its starting position over period of time and it is analogous to cargo’s processivity, that is, the distance that the motors drive the cargo before detachment from microtubules. In general, the larger the MSD, the higher the active transport of the cargo and the higher will be the processivity. MSD was calculated using the following formula, where *τ* is the delay or sliding time, T is the total time, t is the current time point, and x is the position:

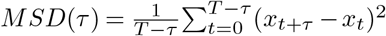

Velocity is the rate of change in position of a cargo in motion from a frame of reference, and since it is a vector quantity, it has a directional component in it. Velocity of the trajectories provides information on the directional switches that the motor proteins undergo when carrying cargoes in a cell. The trajectories which span less than 400 time frames were discarded. Sliding window algorithm was employed in analyzing the data, which reduces the computational power by breaking a large array into smaller sub-arrays. Using this algorithm, the average velocity of individual trajectories was calculated with a window size of 15 seconds. The average of all the trajectories was calculated based on the 20 second period before inhibition, which represents the state of motor motility right before inhibition. Average velocity was then categorized into three types: Positive velocity (velocity*≥*0.01 *µ*m/s), negative velocity (velocity *≤* 0.01 *µ*m/s) and neutral velocity (0.01 *µ*m/s < velocity < -0.01 *µ*m/s).

### Mathematical modeling

Following motor parameters from Gickling et al. (41), with the exception of the binding rates which were reduced 100 fold to match the frequency of directional switches in our experiments, generated cargo trajectories similar to the observed trajectories in our data:

| Parameters | Kin-1 | Kin-2 | Kin-3 | Dyn |
| --- | --- | --- | --- | --- |
| Unbinding rate($\text{s}^{-1}$) | 0.96 | 0.76 | 0.16 | 0.1 |
| Binding rate($\text{s}^{-1}$) | 1.0 | 3.0 | 9.9 | 0.05 |
| Detachment force(pN) | 6.8 | 3 | 1.3 | 100 |
| Stall force(pN) | 6 | 6 | 6 | -3.6 |
| Forward velocity(nm/s) | 586 | 307 | 910 | -360 |
| Backward velocity(nm/s) | 24 | 24 | 24 | -40 |
| No. of motors - EEs | 1 | 0 | 1 | 6 |
| No. of motors - LEs/Lys | 1 | 1 | 1 | 9 |

To simulate different endosome populations, we changed the number of attached motors. For early endosomes, we decreased the number of attached dynein motors based on our reported data (42). Moreover, we omitted kinesin-2 in our model to recapitulate motors reported to be present on early endosomes (31, 58).

### Statistical analysis

Distribution of data was checked on MATLAB using histograms and quantile-quantile plots and data was observed to be not normally distributed. Thus, a Wilcoxon signed-rank test was used to test for statistical significance between dark and lit datasets. A p-value below 0.05 was considered statistically significant.

## Acknowledgements

We would like to thank other students in the lab, namely Abdullah Chaudhary, Linda Balabanian, and Ora Cohen for help with data analysis and thoughtful discussions. We would like to thank the Genome Quebec Innovation Centre at McGill for sequencing our DNA samples. The work was supported by the Canadian Institutes of Health Research (CIHR) grant PJT-159490 to A.G. Hendricks.

## Footnotes

### Author contributions

The project was conceptualised by AGH and SN. SN and AGH designed the experiments. SN and SW cloned the inhibitors. SN performed all the experiments and analysis. KS wrote the code for velocity analysis. SN, FB, and AGH developed the mathematical model. SN and AGH wrote the manuscript with input from SW and KS. AGH obtained funding for this project.

## Supplementary Information

**Fig. S1.**
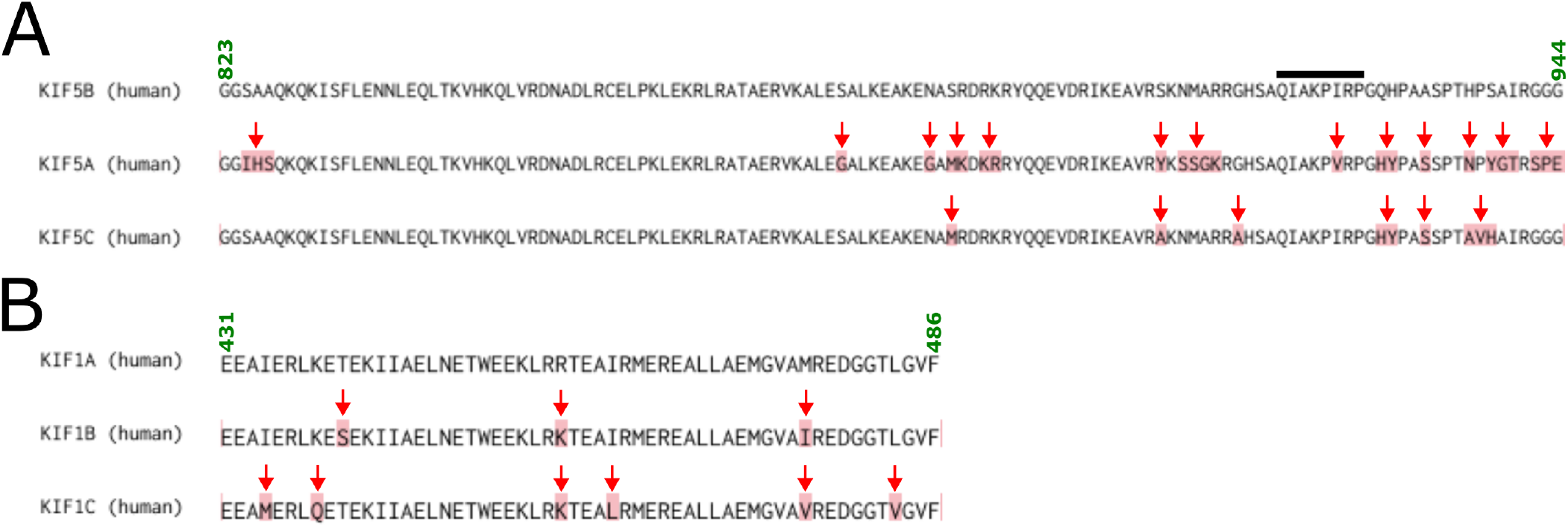
Comparison of amino acid sequences of autoinhibitory domains from different kinesin isoforms. (A) Sequence alignment for inhibitory domain of kinesin-1 (sequence retrieved from UniProt). Black line indicates the sequence responsible for motor inhibition and mismatches are indicated by red arrows. Residue numbers are marked on the top. (B) Sequence alignment for inhibitory domain of kinesin-3 (sequence retrieved from UniProt). Mismatches are indicated by red arrows. Residue numbers are marked on the top.

**Fig. S2.**
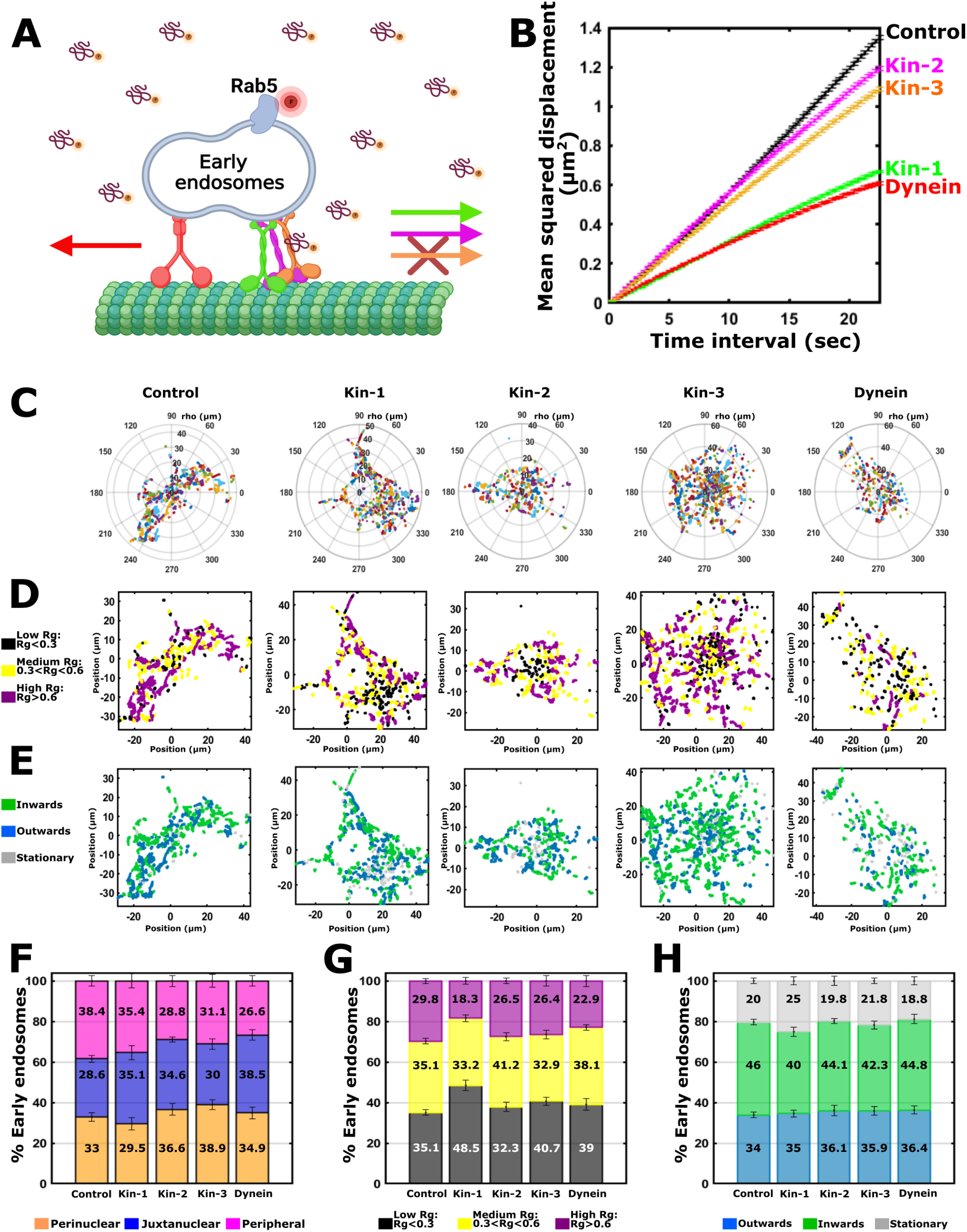
Sustained inhibition of kinesin-1 or dynein motors reduces early endosome motility more notably than inhibiting other motors. (A) Schematic illustration of overexpression studies with early endosomes. (B) Mean squared displacement plot of early endosomes in cells (the mean of all cells) expressing inhibitory peptides for different motor proteins. (C and F) Polar plots of trajectories **Fig. S2**. (shown in various colours) with the cell center as (0, 0). x, y coordinates were used to determine the localisation of cargoes to peripheral, juxtanuclear and perinuclear region of the cell, shown in orange, blue and pink respectively on the stacked bar plot (mean ± SEM) for cells under different inhibitory conditions. (D and G) Radius of gyration thresholds were used to categorize cargo trajectories into low, medium and high Rg trajectories, shown in black, yellow and purple respectively on the cell plot and on the stacked bar plot (mean ± SEM) for cells under different inhibitory conditions. (E and H) The net directionality of cargoes was categorized as outwards, inwards or stationary from the cell center, shown in blue, green and gray respectively on the cell plot and on the stacked bar plot (mean ± SEM) for cells under different inhibitory conditions. COS-7 cells were used for these experiments. Trajectories from the following number of cells were used to prepare plots B, F, G and H :-Control: 4231 trajectories from 22 cells; kinesin-1 inhibition: 2722 trajectories from 13 cells; kinesin-2 inhibition: 3089 trajectories from 12 cells; kinesin-3 inhibition: 2412 trajectories from 11 cells; dynein inhibition: 1601 trajectories from 10 cells.

**Fig. S3.**
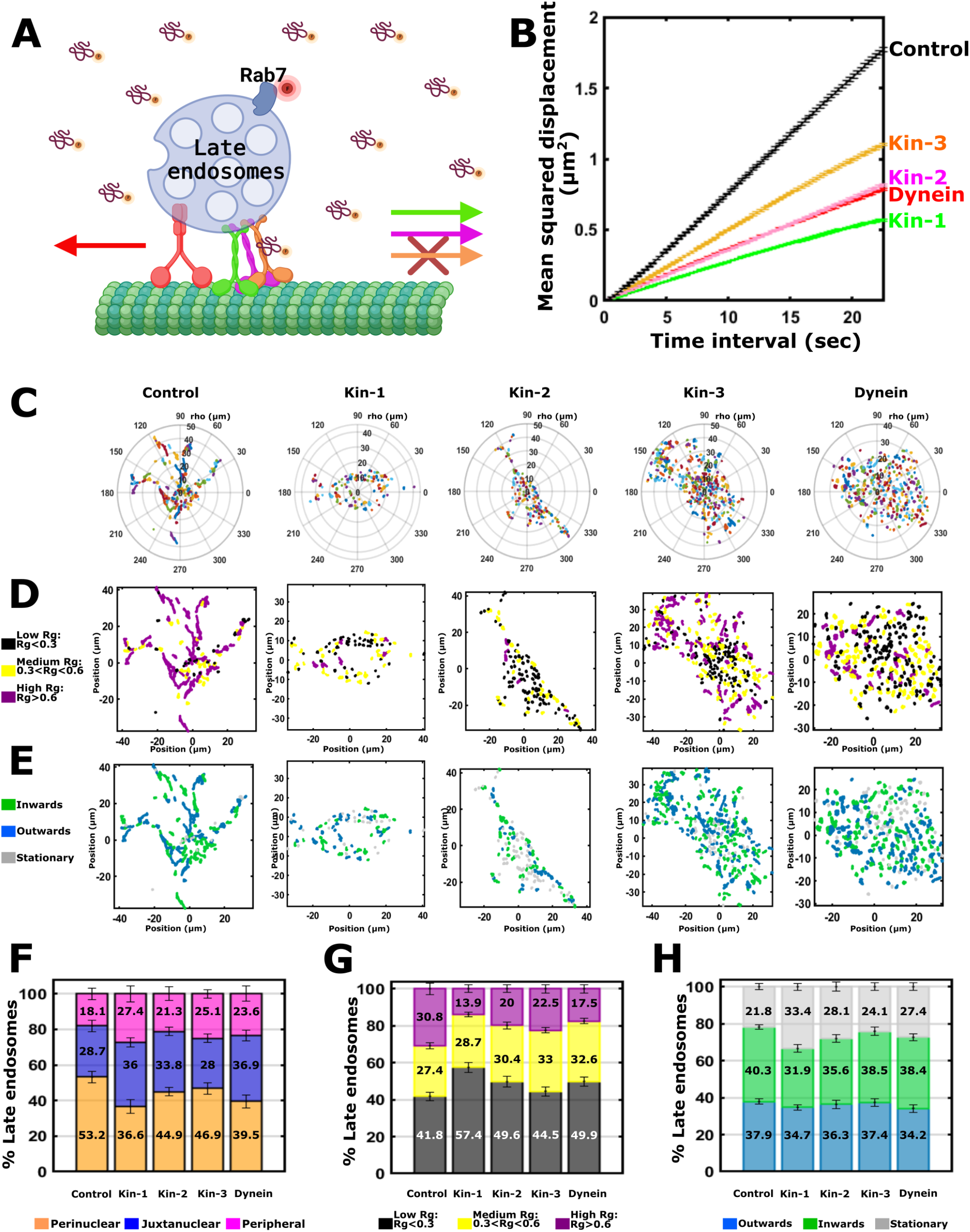
Late endosome motility affected by sustained inhibition of all motors. (A) Schematic illustration of overexpression studies with late endosomes. (B) Mean squared displacement plot of late endosomes in cells (the mean of all cells) expressing inhibitory peptides for different motor proteins. (C and F) Polar plots of trajectories (shown in various colours) with the cell center as (0, 0). x, y Fig. S3. coordinates were used to determine the localisation of cargoes to peripheral, juxtanuclear and perinuclear region of the cell, shown in orange, blue and pink respectively on the stacked bar plot (mean ± SEM) for cells under different inhibitory conditions. (D and G) Radius of gyration thresholds were used to categorize cargo trajectories into low, medium and high Rg trajectories, shown in black, yellow and purple respectively on the cell plot and on the stacked bar plot (mean ± SEM) for cells under different inhibitory conditions. (E and H) The net directionality of cargoes was categorized as outwards, inwards or stationary from the cell center, shown in blue, green and gray respectively on the cell plot and on the stacked bar plot (mean ± SEM) for cells under different inhibitory conditions. COS-7 cells were used for these experiments. Trajectories from the following number of cells were used to prepare plots B, F, G and H :-Control: 1913 trajectories from 11 cells; kinesin-1 inhibition: 2773 trajectories from 17 cells; kinesin-2 inhibition: 2765 trajectories from 12 cells; kinesin-3 inhibition: 3130 trajectories from 13 cells; dynein inhibition: 2263 trajectories from 12 cells.

**Fig. S4.**
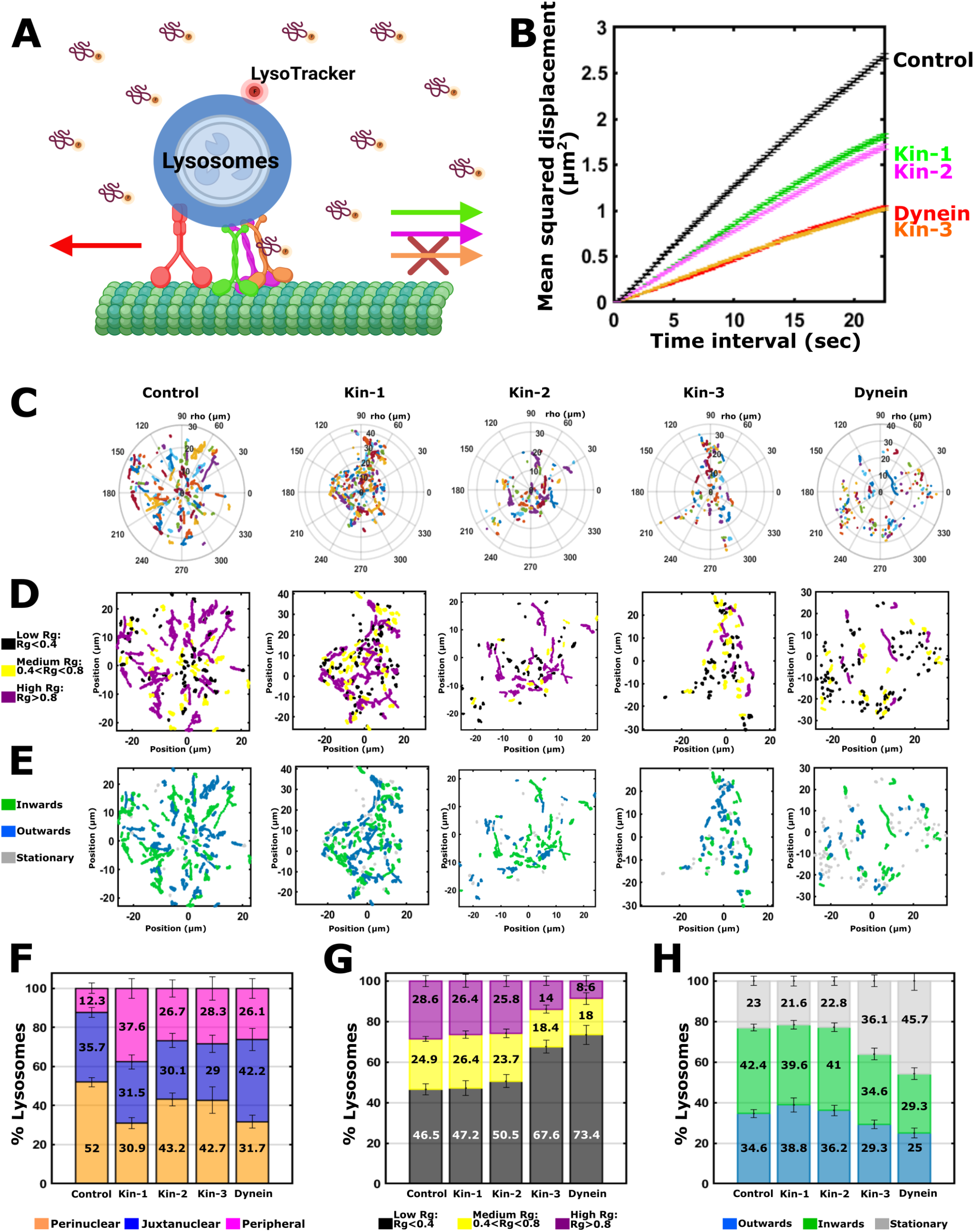
Sustained inhibition of kinesin-3 and dynein shows the strongest reduction in lysosome motility, followed by kinesin-1 and -2 inhibition. (A) Schematic illustration of overexpression studies with lysosomes. (B) Mean squared displacement plot of lysosomes in cells (the mean of all cells) expressing inhibitory peptides for different motor proteins. (C and F) Polar plots of trajectories **Fig. S4**. (shown in various colours) with the cell center as (0,s 0). x, y coordinates were used to determine the localisation of cargoes to peripheral, juxtanuclear and perinuclear region of the cell, shown in orange, blue and pink respectively on the stacked bar plot (mean ± SEM) for cells under different inhibitory conditions. (D and G) Radius of gyration thresholds were used to categorize cargo trajectories into low, medium and high Rg trajectories, shown in black, yellow and purple respectively on the cell plot and on the stacked bar plot (mean ± SEM) for cells under different inhibitory conditions. (E and H) The net directionality of cargoes was categorized as outwards, inwards or stationary from the cell center, shown in blue, green and gray respectively on the cell plot and on the stacked bar plot (mean ± SEM) for cells under different inhibitory conditions. COS-7 cells were used for these experiments. Trajectories from the following number of cells were used to prepare plots B, F, G and H :-Control: 1345 trajectories from 13 cells; kinesin-1 inhibition: 1417 trajectories from 10 cells; kinesin-2 inhibition: 950 trajectories from 11 cells; kinesin-3 inhibition: 883 trajectories from 16 cells; dynein inhibition: 707 trajectories from 12 cells.

**Fig. S5.**
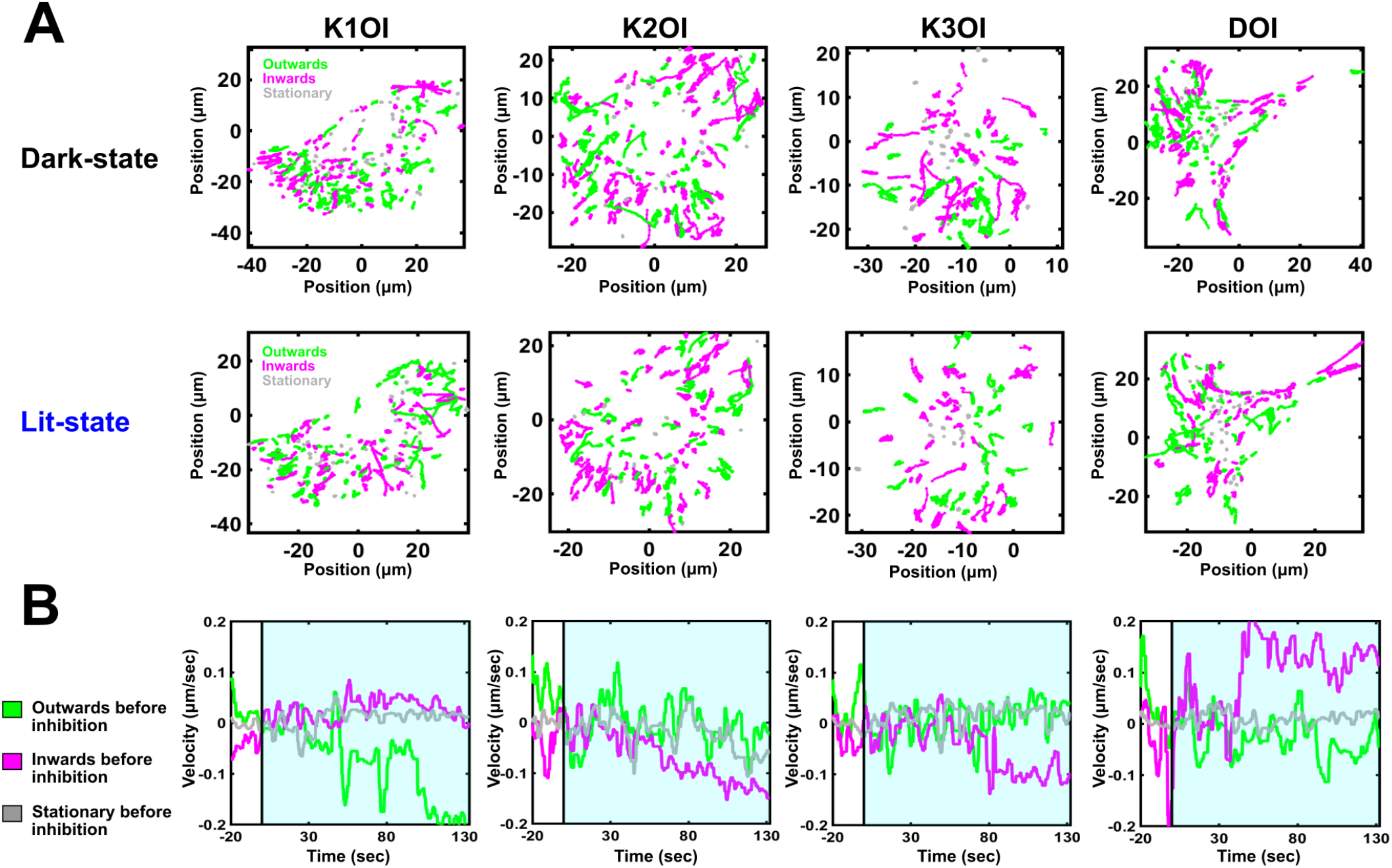
Analysis of the velocity for additional cells shows the change in lysosome transport upon optogenetic inhibition. (A) Polar plot projections of lysosomes trajectories from time-lapse images, centered around the cell nucleus, showing their directionality in a U2OS cell under dark-state (upper panel) and lit-state (bottom panel) conditions. The four panels correspond to cells that were transiently transfected with different optogenetic inhibitors. The net directionality was categorized as inwards (magenta), outwards (green) or stationary (gray) based on Rg values, and rho values in the first and the last points of the trajectories. (B) Plot shows the changes in average velocity for all the trajectories in a cell (corresponding to the cell shown in A) upon blue light illumination. For velocity analysis, average velocity was first categorised into three types, namely, positive velocity, negative velocity, and neutral velocity. It was then normalized to the average velocity in the time window just before inhibition, allowing us to compare changes at the time of inhibition. The color scheme is also based on the average velocity of the trajectories right before the inhibition, where green represents positive average velocity before inhibition, magenta represents negative average velocity before inhibition, and gray represents stationary vesicles that were not moving before inhibition.

**Fig. S6.**
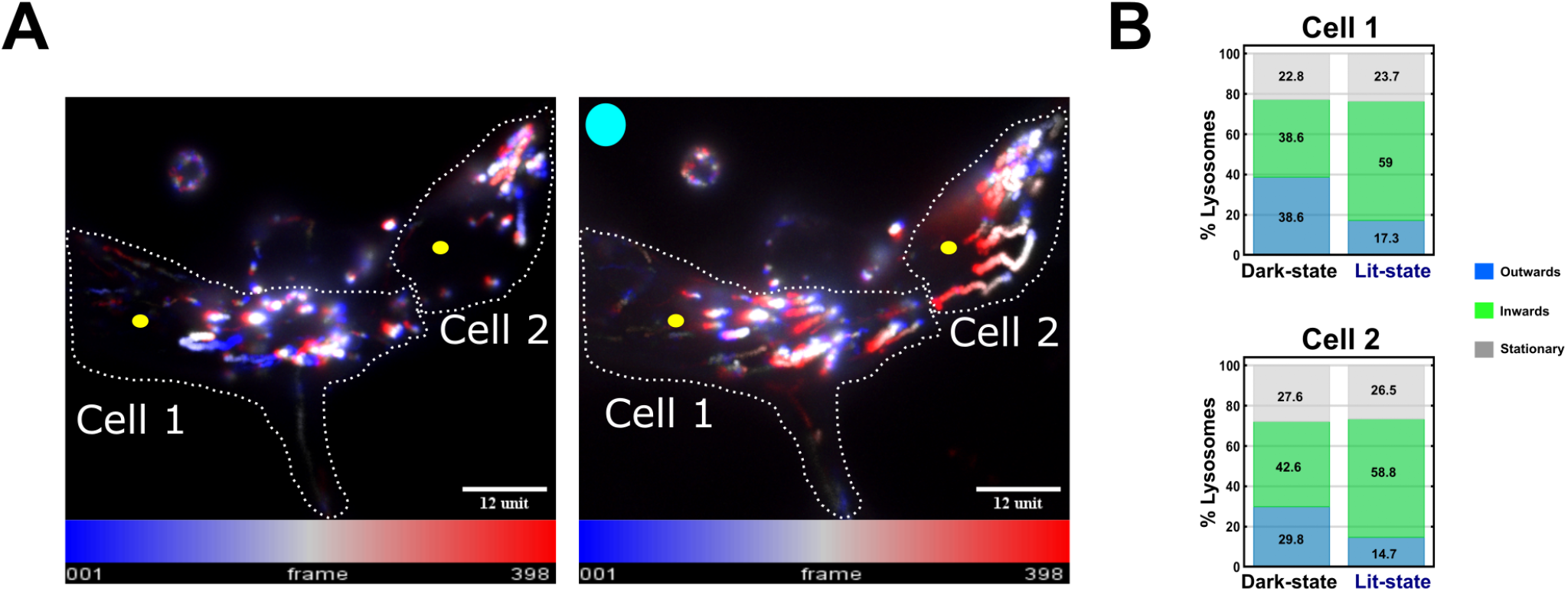
Combinatorial inhibition of multiple kinesins drastically steers the directionality of cargoes to retrograde direction. (A) Temporal maximum projection images of lysosome motility in U2OS cells co-expressing optogenetic kinesin-2 and kinesin-3 inhibitors in the dark state (on the left) and in the lit-state (on the right). Cell center is shown by yellow dot. (B) Stacked bar plots showing net directionality of cargoes categorized as outwards, inwards or stationary from the cell center, shown in blue, green, and gray respectively for cell 1 (on top) and cell 2 (below).

**Fig. S7.**
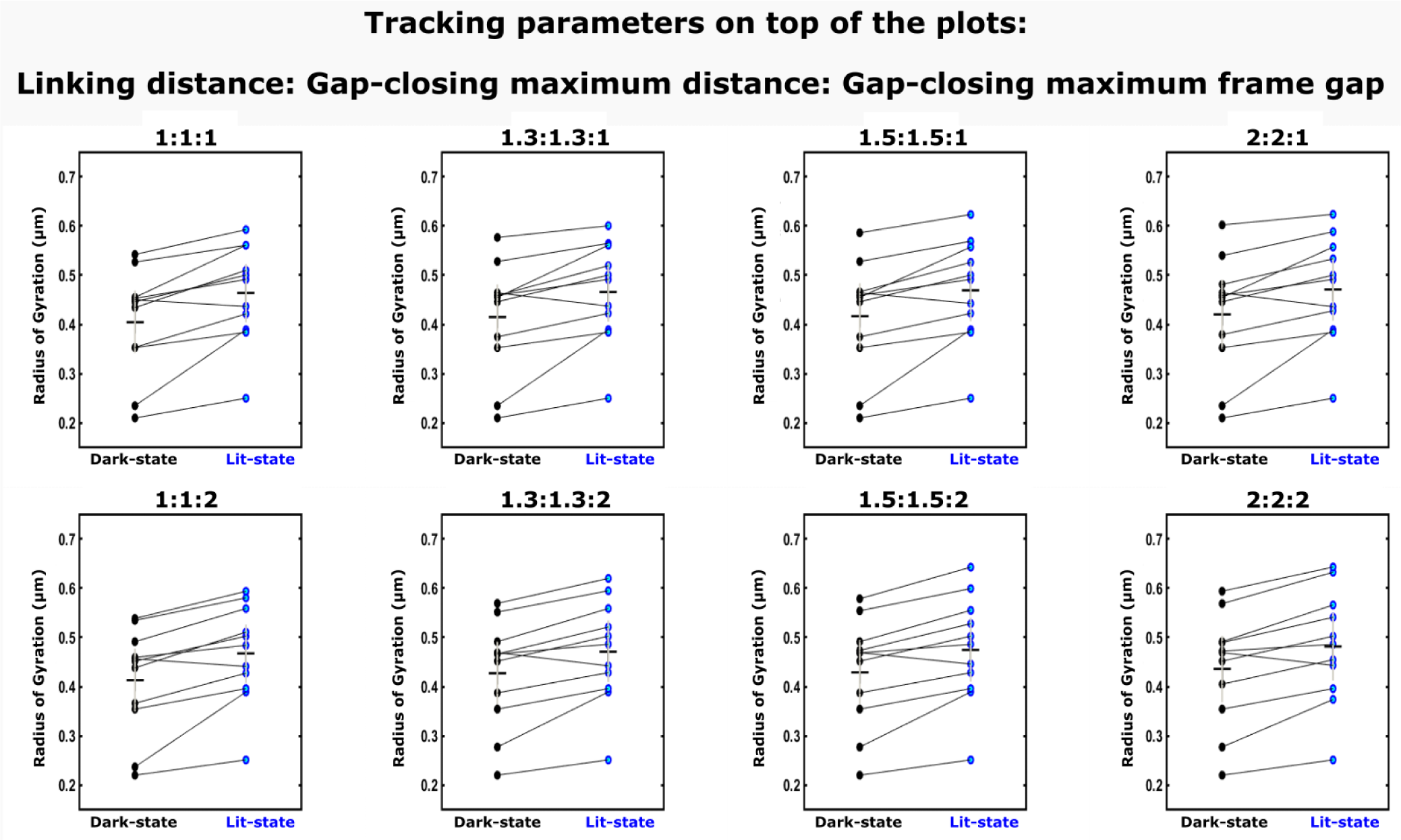
Characterisation of tracking parameters. Radius of gyration plots for a subset of data taken from dynein inhibition of Rab5 vesicles. Cargo motility in this dataset was tracked using different tracking parameters that are available on TrackMate. The tracking parameters are indicated on top of each plot in the format linking distance: gap-closing maximum distance: gap-closing maximum frame gap.

**Supplementary video**. Optogenetic cell shown in Figure 1D, demonstrating the release of mCherry-tagged inhibitor from the mitochondria upon 10s blue light pulse at 10s, 45s and 85s in the video.

